# Aftiphilin regulation of myosin light chain kinase activity promotes actin dynamics and intestinal epithelial barrier function

**DOI:** 10.1101/2022.03.15.484511

**Authors:** Ivy Ka Man Law, Kai Fang, Charalabos Pothoulakis, Carl Robert Rankin

## Abstract

The expression levels of aftiphilin (AFTPH) are significantly lower in inflamed colonic tissues from patients with ulcerative colitis (UC) and mice with experimental colitis. During colonic inflammation, the selective permeability of the colonic epithelium is compromised largely due to dysregulation of proteins associated with either the tight junction (TJ) complex and actin-myosin contraction rings. Here, we hypothesized that inflammation-associated reduction in AFTPH levels might cause an increase in the selective permeability of the colonic epithelium. In this study, we measured the transepithelial electric resistance (TEER), sodium (Na^+^) ion flux and dextran permeability in polarized colonic epithelial cells after manipulation of AFTPH. Silencing of AFTPH reduced TEER, increased Na^+^ ion flow and dextran permeability. Examination of mRNA and protein levels of multiple TJ proteins and Na^+^ ion transporters suggested that AFTPH deficiency did not significantly change expression of most of these transmembrane proteins. While the gross structure of the TJs in AFTPH gene-silenced cells appeared normal, elevated levels of junctional Occludin were observed. Most notably we observed that AFTPH co-localized with myosin light chain kinase (MLCK) and attenuated cellular MLCK activity as observed by phospho-myosin light chain 2 (pMLC2) western blots. Importantly, inhibition of MLCK activity reversed the reduction of TEER in AFTPH-deficient monolayers. Lastly, examination on transmission electron microcopy on microvilli and immunofluorescent microscopy on actin filament arrangement showed that AFTPH deficiency also affected filament arrangement in colonic epithelial cells. Taken together, these results suggest that AFTPH regulates intestinal epithelial permeability and actin polymerization in colonic epithelium through interfering MLCK/MLC interactions.

## Introduction

The intestinal barrier is formed by a single layer of polarized intestinal epithelial cells located between the underlying mucosa and lumen. At homeostasis, intestinal epithelial cells maintain a selective barrier, allowing the paracellular flux of ions, nutrients and water [review in (1)], through dynamic regulations of (i) tight junction (TJ) transmembrane proteins; (ii) contraction of actin-myosin rings; and (iii) ion transporters. TJ transmembrane proteins from adjacent epithelial cells create a functional gate to the paracellular flux of small ions and water (also known as the pore pathway) at the boundary between the apical and basolateral domains of intestinal epithelium. TJ transmembrane proteins can be grouped into three main families, claudins, TJ-associated MARVEL proteins (TAMPs), and the cortical thymocyte marker in *Xenopus* (CTX). Claudins are important in regulating paracellular, size-dependent, flux of charged ions and can be further divided into two groups, the sealing claudins and the pore-forming claudins [review in (2)]. For example, claudin-1 (CLDN1) is one of the sealing claudins important for maintaining structure (3) and is translocated and downregulated during proinflammatory cytokine-(4) and hypoxia-(5) induced barrier dysfunction. In contrast, claudin 2 (CLDN2), a member of pore-forming claudins, is responsible for creating water and sodium (Na^+^) ion channels across the polarized epithelium (6,7) and its loss is associated with attenuated colitis development (8). Furthermore, members of the claudin family interact with TAMPs and CTX in order to maintain the structural integrity of TJs (9,10), while individual TAMPS and CTX also act as anchors to various signaling molecules [review in (11)]. Overall, the TJ transmembrane proteins work together to regulate the passive transport of water and small ions, provide structural integrity to TJ and enhance signaling cascades.

Myosin light chain (MLC) is part of non-muscle myosin (NMM) heterohexamer which forms the actin-myosin ring present at the cell periphery and linked to the TJ protein complex [review in (12,13)]. In general, contraction of actin-myosin rings in individual intestinal epithelial cells contributes to widening of intercellular spaces and subsequently allowing macromolecules to passively pass through paracellular space of the intestinal epithelium (leak pathway), and thus compromising intestinal barrier function. During intestinal inflammation, stimulation from proinflammatory cytokines on intestinal epithelium activates myosin light chain kinase (MLCK) and promotes phosphorylation of MLC2 (14), which leads to contraction of actin-myosin ring (15) and reduced actin anchoring to TJ protein complex (16), resulting in a reduction in intestinal epithelial barrier function.

Lastly, in addition to the passive transport of ions, water and macromolecules through paracellular space, charged ions and macromolecules are also actively transported through epithelial cells from lumen through various ion transporters present in both apical and basolateral sides of intestinal epithelial cells. Intestinal epithelial Na^+^ ion transporters, such as, the sodium-hydrogen exchanger / solute carrier family 9 member 3 (SLC9A3), Na^+^-glucose cotransporter and Na^+^/K^+^ transporting ATPase, contribute to the active transport of Na^+^ ions across colonic epithelium [review in (17)]. Coincidently, activities of SLC9A3 and Na+-glucose cotransporter have been shown to affect electrical potential and MLC activation across intestinal epithelial cells (18,19). Taken together, the flow of ions, water and nutrients across paracellular space is governed by intricate expression and interactions between TJ transmembrane proteins, MLC phosphorylation and ions transporters present on intestinal epithelial cells.

In patients with Inflammatory Bowel Diseases (IBD), a compromised barrier is often associated with recurrent and relapsing intestinal inflammation [review in (1,20)]. Our previous studies have identified aftiphilin (AFTPH) to be downregulated during active colonic inflammation in colonic tissues from patients with ulcerative colitis (UC) and experimental colitis mouse models (21). Since previous studies have suggested that AFTPH is a binding partner of non-muscle myosin 2 (NMMII) (22,23), we hypothesized that AFTPH might interact with components regulating epithelial permeability in colonic epithelial cells. In this study, we examined transepithelial electrical resistance (TEER), Na+ ion flux, expression of TJ transmembrane protein, MLCK activity and actin dynamics in intestinal epithelial cells in the presence and absence of AFTPH and showed that AFTPH deficiency increased epithelial cell permeability potentially by increasing MLCK activity and interfering with actin polymerization.

## Results

### Aftiphilin downregulation compromises intestinal epithelial barrier function in vitro

According to The Human Protein Atlas project (https://www.proteinatlas.org/ENSG00000119844-AFTPH/cell) and other previous studies (23-26), AFTPH is expressed in most of the major cell types in the gut, such as, endothelial cells, immune cells, fibroblasts and neurons, in addition to our findings on colonic epithelial cells (21). Since compromised intestinal epithelial permeability is a hallmark of ulcerative colitis, here, we hypothesize that reduced AFTPH expression has a direct role in increasing paracellular permeability of colonic epithelial monolayer. To accomplish this, we transduced three different colonic epithelial cell lines with shRNA containing lentiviruses: NCM460, Caco2-BBe & T84. NCM460 cells are non-transformed colonic epithelial cells derived from normal human colonic mucosa (27). Caco2-BBe and T84 cells are both intestinal epithelial cell lines derived from colorectal adenocarcinoma and differentiated into epithelial monolayers, closely resembling human colonic epithelium (28). As shown in Fig.1, NCM460, Caco2-BBe and T84 cell lines transduced with the AFTPH-targeting shRNA have 51.3±7.29%, 96.0±0.51% and 93.6±3.29% reduction in AFTPH protein levels, when compared to same cell lines transduced with scramble shRNA, respectively (*p*=0.0385, *p*=0.0072, *p*=0.0492). We then tested the formation and maintenance of permeability, which can be represented by TEER, of AFTPH gene-silenced Caco2-BBe and T84 monolayers. As shown in Fig. 2A, AFTPH-deficient Caco2-BBe developed significantly lower TEER, while AFTPH deficiency significantly delayed the development of TEER in T84 monolayers. Since TEER is a measurement of overall ion flow through intestinal epithelium *in vitro*, we next examined Na^+^ ion and dextran permeability in Caco2-BBe and T84 monolayers gene-silenced with AFTPH and their control counterparts, as a representation of the flux of small, charged ions and uncharged molecules. Our results showed that, in both Caco2-BBe and T84 cells, AFTPH gene-silencing not only significantly lowered TEER (Fig. 2A), but the dilution potential of Na^+^ ions was also lowered (12.3±0.25 mV vs. 10.9±0.15 mV; and 14.6±0.64 mV vs. 11.4±0.32 mV, *p*=0.0003), when compared to control cells (Fig. 2B). Interestingly, with regards to flux of 3000, 10,000 and 40,000 M.W. dextran passing across the epithelial monolayers of both cell lines, our results showed differences between Caco2-BBE and T84 gene-silenced cells. In Caco2-BBe cells, AFTPH gene-silencing did not show altered permeability to dextran of all molecular weights (Fig. 2C-E). In contrast, AFTPH-deficient T84 monolayers (Fig. 2D-F) showed increased permeability to dextran of 3,000 M.W. (2.4±0.38 pmol/cm^2^/h vs. 4.2±0.44 pmol/cm^2^/h, *p*<0.0001), 10,000 M.W. (0.3±0.09 pmol/cm^2^/h vs. 1.0±0.19 pmol/cm^2^/h, *p*=0.02) and 40,000 M.W (0.006±0.0014 pmol/cm^2^/h vs. 0.009±0.0022 pmol/cm^2^/h, *p*=0.027). Taken together, AFTPH deficiency increased epithelial permeability, potentially by increasing permeability to Na^+^ ions and, to a lesser extent, large molecules in colonic epithelial monolayer.

**Fig. 1.**
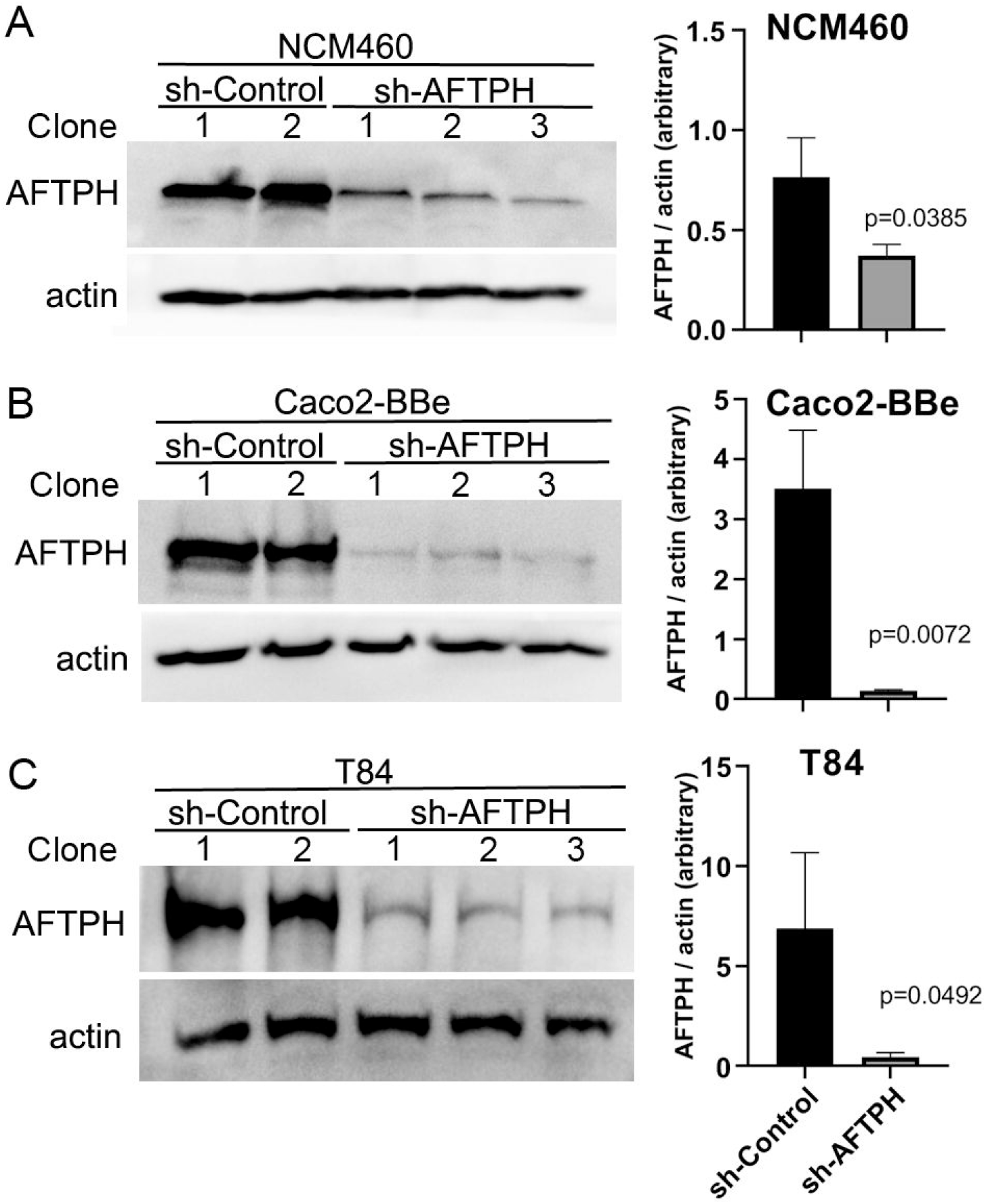
AFTPH expression is significantly reduced in colonic epithelial cell lines transduced with recombinant lentivirus expressing sh-AFTPH. Control recombinant lentivirus and recombinant lentivirus carrying sh-AFTPH were used to transduced 3 cell lines, (A) NCM460 (sh-control = 2, sh-AFTPH = 3), (B) Caco2-BBe (control = 2, sh-AFTPH = 3) and (C) T84, transduced with recombinant lentivirus expressing sh-AFTPH (control = 2, sh-AFTPH = 3). Individual clones were generated from sh-control and sh-AFTPH transduction. Expression of AFTPH in different clones was examined using Western Blot and densitometric analysis. Mean± SEM.

**Fig. 2.**
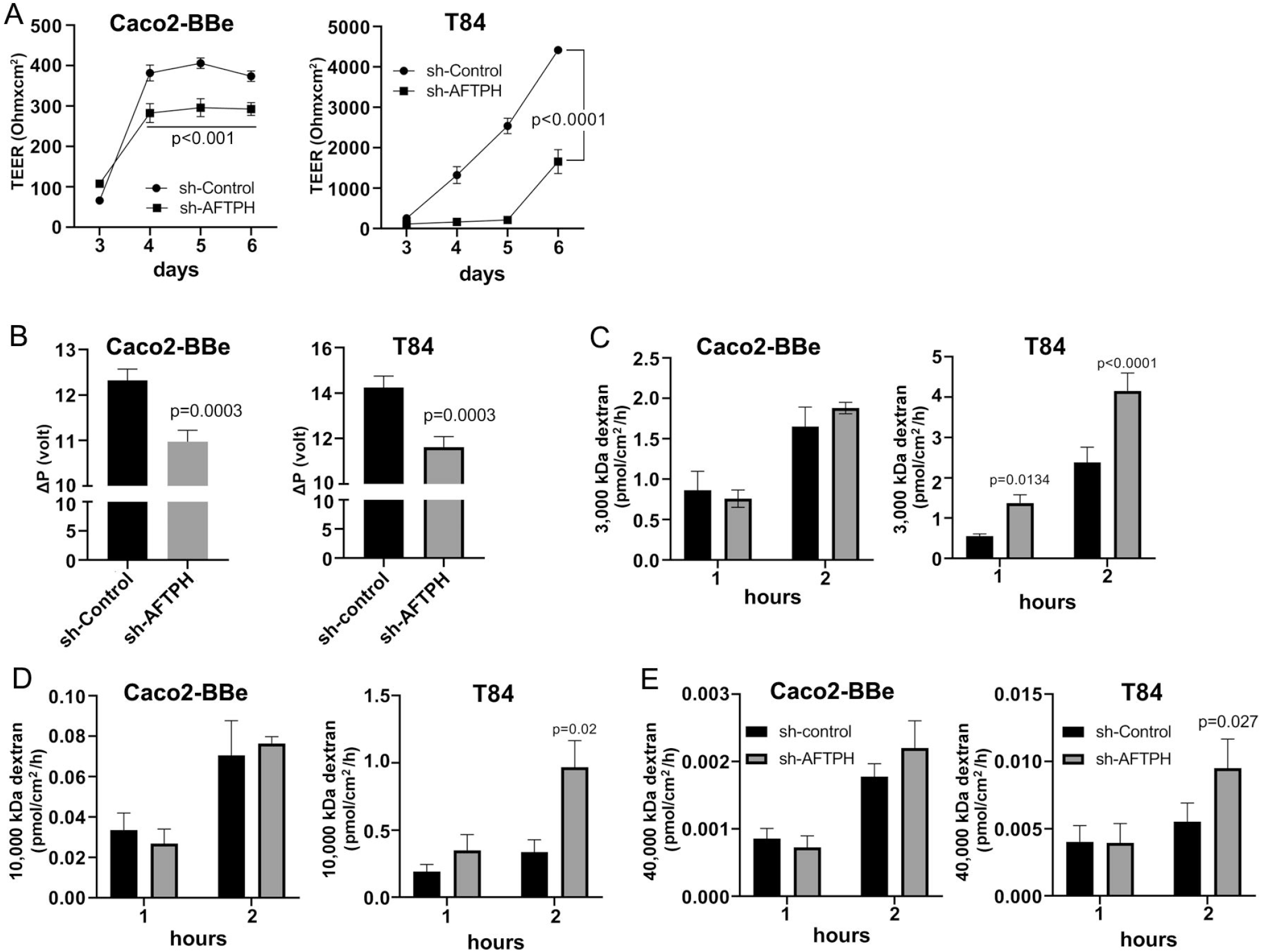
AFTPH gene-silencing significantly increases colonic epithelial permeability in Caco2-Bbe and T84 monolayers. (A) Transduced sh-control and sh-AFTPH clones were plated on transwells on day 0 and allowed to polarize. A volt-ohm meter was used to measure Transepithelial electrical resistance (TEER) on each day from 3 to 6 after plating. Mean± SEM. (B) Na^+^ ion permeability was measured by dilution potential assay as described in the Methods at day 6. Mean± SEM. (n=3). Dextran of size (C) 3000, (D) 10,000, (E) 40,000 M.W. was incubated with polarized sh-control and sh-AFTPH Cac2-BBe and T84 clones in dextran permeability assay as described in Methods at day 6 and measured at 1 and 2 hours after the start of the experiment. Mean± SEM. (n=3).

### Occludin expression is modulated by AFTPH gene-silencing in intestinal epithelial monolayers

Since Na^+^ ion flow is commonly elevated in both AFTPH-deficient cell lines, we next tested the role for AFTPH in the two major mechanisms for regulating paracellular ion flow, active ion transport (through transcellular Na^+^ ion transporters) and passive ion flow (pore pathway, through TJ proteins). We first examined gene expression of multiple claudins and OCLN and discovered that AFTPH gene-silencing increased gene expression of claudin-1 (CLDN1) in both Caco2-BBe and T84 monolayers by 1.6±0.08 fold (*p*=0.0259) and 11.2±3.68 fold (*p*=0.0334), respectively (Fig. 3A). Expression levels of claudin-2, -3 and -4 (CLDN2, CLDN3, CLDN4) were also altered in both Caco2-BBe and T84 monolayers. Specifically, significantly higher expression levels of CLDN2 and lower expression levels of CLDN3 were observed in AFTPH-deficient Caco2-BBe cells (*p*=0.004, *p*=0.0252, respectively). However, CLDN2 and CLDN3 mRNA levels were not altered in sh-AFPTH-transduced T84 monolayers (Fig. 3B&C). Alternately, CLDN4 expression was significantly upregulated in T84 monolayers stably transduced with sh-AFTPH (*p*=0.0208, Fig. 3D), but, in Caco2-BBe cells, AFTPH gene-silencing did not affect CLDN4 expression. Interestingly, AFTPH deficiency slightly increased OCLN mRNA in both Caco2-BBe and T84 monolayers, although no statistical significance was reached (Fig. 3E). Next, we examined expression of Na^+^ ion membrane transporters. Our results showed that SLC9A3 expression was significantly increased only in Caco2-BBe cells (*p*=0.0103, Fig. 3F); and expression of Na^+^/K^+^ transporting ATPase subunit ATP1B1 was significantly upregulated in T84 cells but not Caco-2 cells (*p*=0.0179, Fig. 3H). ATP1A1, another Na^+^/K^+^ ATPase subunit, was not significantly altered in either cell line (Fig. 3G). Overall, our gene expression analysis demonstrated that reduced AFTPH expression did not affect the overall gene expression of these genes except CLDN1 and OCLN in intestinal epithelial cells.

**Fig. 3.**
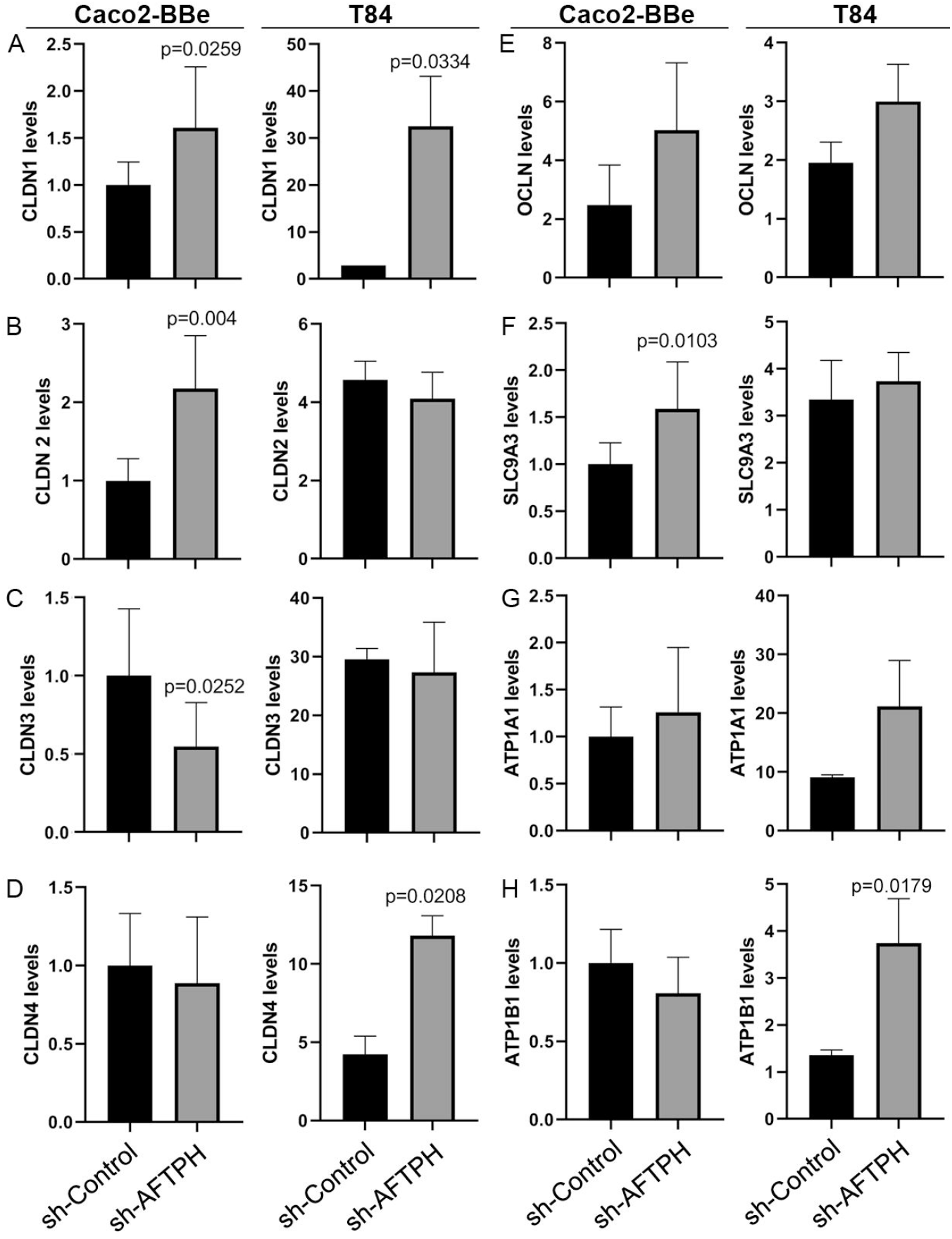
Reduced AFTPH expression does not significantly alter mRNA levels of tight junction proteins and Na^+^ ion transporters in Caco2-BBe and T84 monolayers. After extracting RNA from polarized cells, RT-qPCR was performed to determine the gene expression of (A-D) CLDN1, -2, -3, -4; (E) OCLN, (F) SLC9A3, (G) ATP1A1 and (H) ATP1B1 in Caco2-BBE and T84 cells using qPCR. Mean± SEM. (n=3).

To determine whether the relative mRNA levels of TJ proteins correlate with protein alterations, protein expression of CLDN1 and OCLN in Caco2-BBe and T84 monolayers were further examined by western blot. Contrary to the results by qPCR, protein expression of CLDN1 was not significantly increased in both Caco2-BBe and T84 monolayers (Fig. 4A&B). Similar to results from gene expression analysis, OCLN expression was slightly, but not significantly, increased in both cell lines (Fig. 4A&B).

**Fig. 4.**
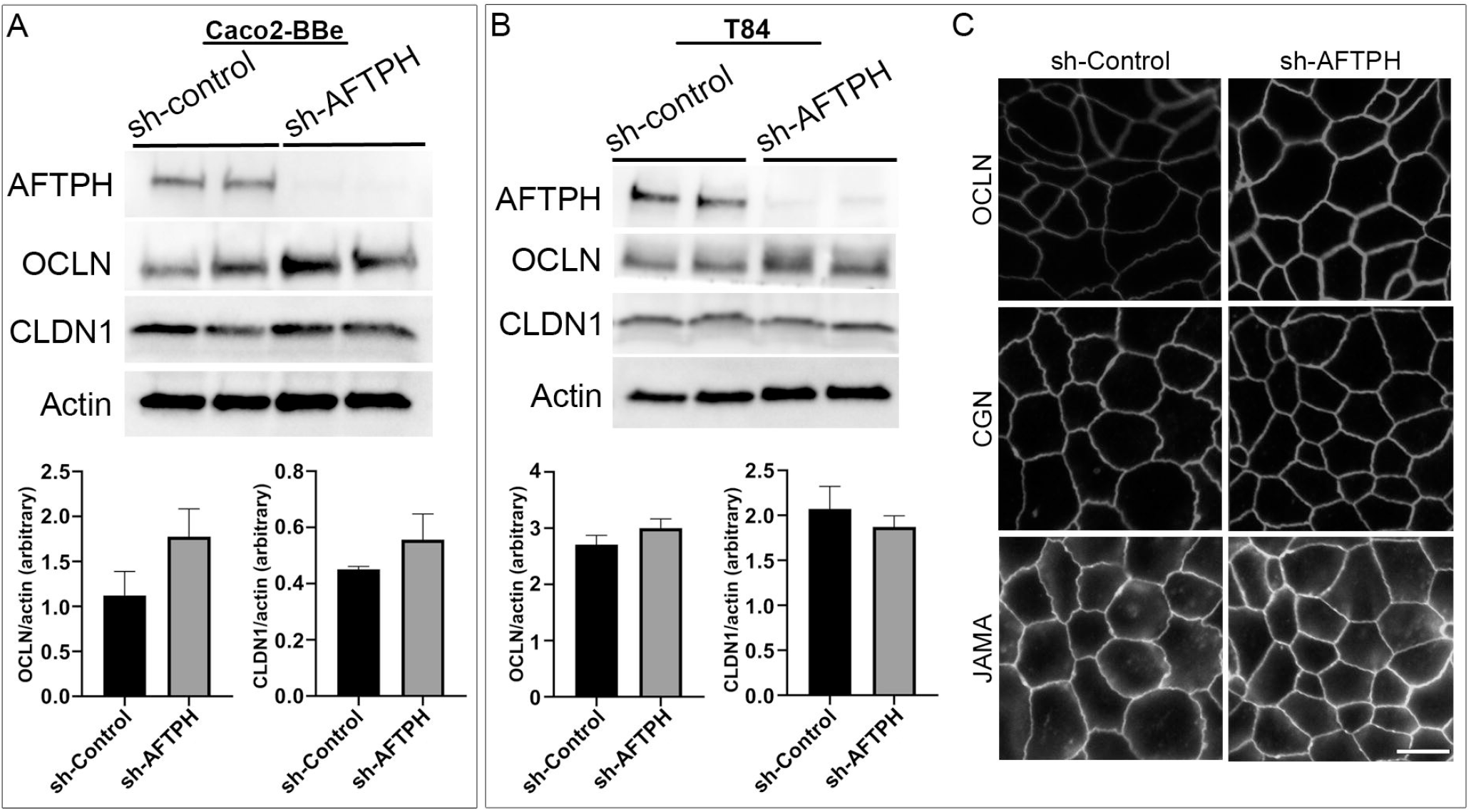
Reduced AFTPH expression increases Occludin membrane localization but does not significantly alter the structure of tight junction in T84 monolayers. (A) Caco2-BBe and (B) T84 sh-control and sh-AFTPH clones (n=2) were seeded and cultured in 6-well plate and harvested. Protein expression of AFTPH, OCLN, CLDN1 were examined using Western Blot, and compared as actin (loading control). Mean± SEM. (C) Structure of tight junction was visualized with antibodies against OCLN, CGN and JAMA using confocal microscopy in T84 cells. Time-exposure used in T84 monolayers transduced with sh-Control- and sh-AFTPH was specific to the antibodies used in labelling. This experiment was repeated 3 times. (Scale = 10 μm).

Another factor leading to increase in intestinal epithelial permeability is translocation of TJ proteins from the plasma membrane to the cytosol. To examine the plasma membrane localization of TJ and its associated proteins, immunofluorescence against OCLN, junctional adhesion molecule-A (JAM-A, a member of the CTX family) and cingulin (CGN) in acetone-fixed T84 monolayers was performed. OCLN and JAM-A are transmembrane proteins that form part of the TJ protein complex (9,29,30), while CGN is a linker protein connecting TJ protein complex and cytoskeleton (31-33) in intestinal epithelial cells. Results from immunofluorescence microscopy of T84 monolayers showed that AFTPH deficiency notably increased OCLN localization on the plasma membrane (Fig. 4C). However, levels of plasma membrane-localized CGN and JAM-A remain similar in monolayers transduced with sh-control and sh-AFTPH (Fig. 4C). These results suggest that enhanced permeability due to AFTPH deficiency is unlikely due to changes in the levels of TJ proteins and their localization but could be explained by stronger OCLN membrane localization.

### AFTPH colocalizes with myosin light chain kinase and regulates its activity in intestinal epithelial monolayers

As we observed that AFTPH regulated dextran flux in colonic epithelial T84 monolayers (Fig. 2), we hypothesized that AFTPH deficiency promotes MLCK activation, promoting the phosphorylation of MLC2 at Ser 19, which would subsequently lead to contraction of peri-junctional actin-myosin rings and disrupted barrier function (15). Interestingly, inactivation of MLCK activity has been shown to increase TEER of intestinal epithelial monolayers (15,34). To test this hypothesis, we examined levels of MLC2 phosphorylation by western blot in individual sh-AFTPH-transduced T84 clones and their control counterparts and demonstrate that levels of pMLC2 were higher in AFTPH-gene silenced T84 cells by 8.8±1.26 fold (*p*=0.0043, Fig. 5A), when compared to control cells, demonstrating that AFTPH deficiency promotes MLCK activity and MLC activation in colonic epithelial cells. To determine if MLCK activity was required for the increased permeability in loss of function AFTPH cells, we blocked MLCK activity in T84 AFTPH-deficient and control cells by treating the cells with PIK, a pharmacological inhibitor of MLCK (35). Results from the measurement of TEER demonstrated that treatment of PIK reduced permeability of AFTPH-deficient T84 monolayers to levels similar to that of the control cells (Fig. 5B), suggesting that increased overall permeability of sh-AFTPH-transduced T84 monolayers was partly due to elevated MLCK activity in these cells. To explore a potential mechanism leading to AFTPH interacting with MLCK in colonic epithelial cells, localization of AFTPH and MLCK were visualized by immunofluorescence confocal microscopy in control T84 and Caco2-BBe monolayers. Interestingly, a high-level of colocalization of AFTPH and MLCK was observed (Fig. 5C), suggesting that some of MLCK molecules were localized at intracellular vesicles or organelles (25). In contrast to control polarized epithelia, AFTPH-deficiency in both cell lines greatly reduced intracellular MLCK localization, despite the increased MLCK activity as shown in Fig. 5A&B. In summary, our results suggest that AFTPH regulates colonic epithelial permeability partly through regulating MLCK localization and activity.

**Fig. 5.**
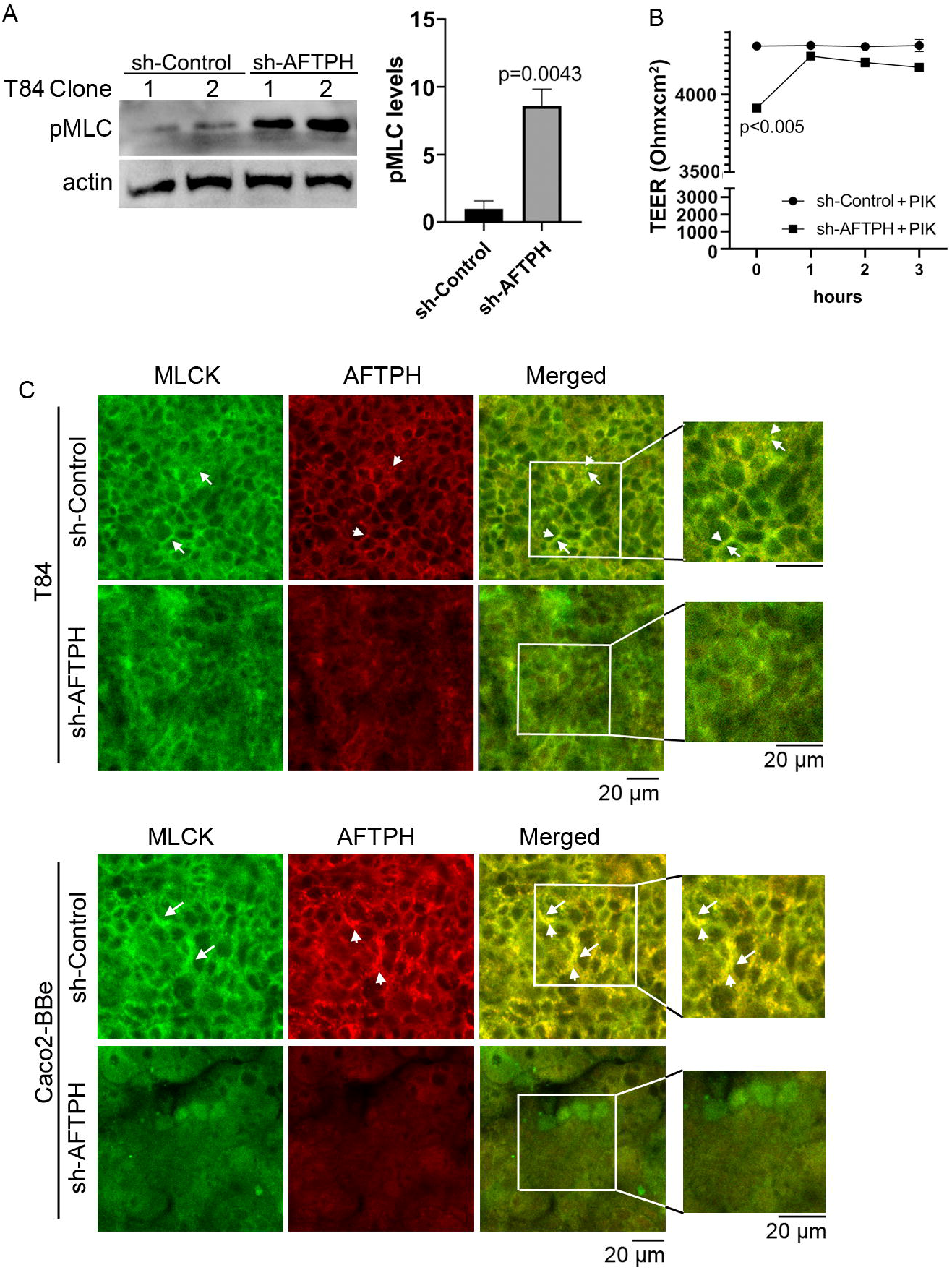
AFTPH colocalizes with MLCK and its deficiency increases MLCK activity. (A) Individual T84 sh-control (n=2) and sh-AFTPH (n=3) clones were cultured in 6-well plates and were harvested when confluent. Levels of phosphorylated MLC2 in different T84 clones was examined by Western blot. Mean ± SEM. (B) Polarized T84 cells were treated with 5 μM PIK at day 6 and TEER was measured every hour (n=3). (C) Polarized T84 and Caco2-BBe cells were cultured for 6 days and colocalization of MLCK (arrow) and AFTPH (arrowhead) was examined by immunofluorescence microscopy. This experiment was repeated 3 times. (Scale = 20 μm).

### AFTPH deficiency dysregulates actin assembly and increases microvilli length at brush border

Since results from this study suggested that AFTPH may regulate activities of MLCK, which has two actin-binding domains, and also closely interact with non-muscle myosin (NMM) (22,23), an actin-based motor protein; we next examined actin assembly at brush borders and the cell periphery by fluorescence microscopy after AFTPH deficiency. Filamentous (F-) actin in non-transformed NCM460 cells at brush borders was significantly dysregulated upon AFTPH depletion, although F-actin at the cell periphery was not altered by the absence of AFTPH (Fig. 6A). We then labelled F-actin and G-actin individually in control and AFTPH-deficient NMC460 and T84 cells and measured overall F-/G-actin ratio. Results from fluorescence spectrometry revealed that AFTPH deficiency significantly lowered F/G-actin ratio in NCM460 and T84 cells (p=0.0259, p<0.0001, respectively), while the individual measurements of F- and G-actin were not different in the presence or absence of AFTPH (Fig. 6B). In addition, confocal microscopy of T84 monolayers also showed no differences in peri-junctional actin-myosin rings under AFTPH deficiency (Fig. 6B). Lastly, we also studied the morphology of microvilli in control and AFTPH-deficient polarized T84 epithelia. T84 monolayers were fixed and imaged by transmission electron microscopy and interestingly, we found that microvilli were more abundant and longer on cell surface of AFTPH-deficient T84 cells (Fig. 6C, as indicated by arrows). A close look at the electron micrographs suggested that F-actin cores in terminal webs of AFTPH-deficient T84 monolayers were also longer than those in control monolayers (Fig. 6C, as indicated by arrowheads). Thus, AFTPH may directly or indirectly regulate actin polymerization, affecting the structure of F-actin in microvilli in intestinal epithelial cells. Taken together, results from our study suggested that AFTPH colocalized with MLCK and regulated its activity on MLC and actin, subsequently affecting intestinal barrier function and microvilli growth, without affecting TJ protein expression, gross structure and localization.

**Fig. 6.**
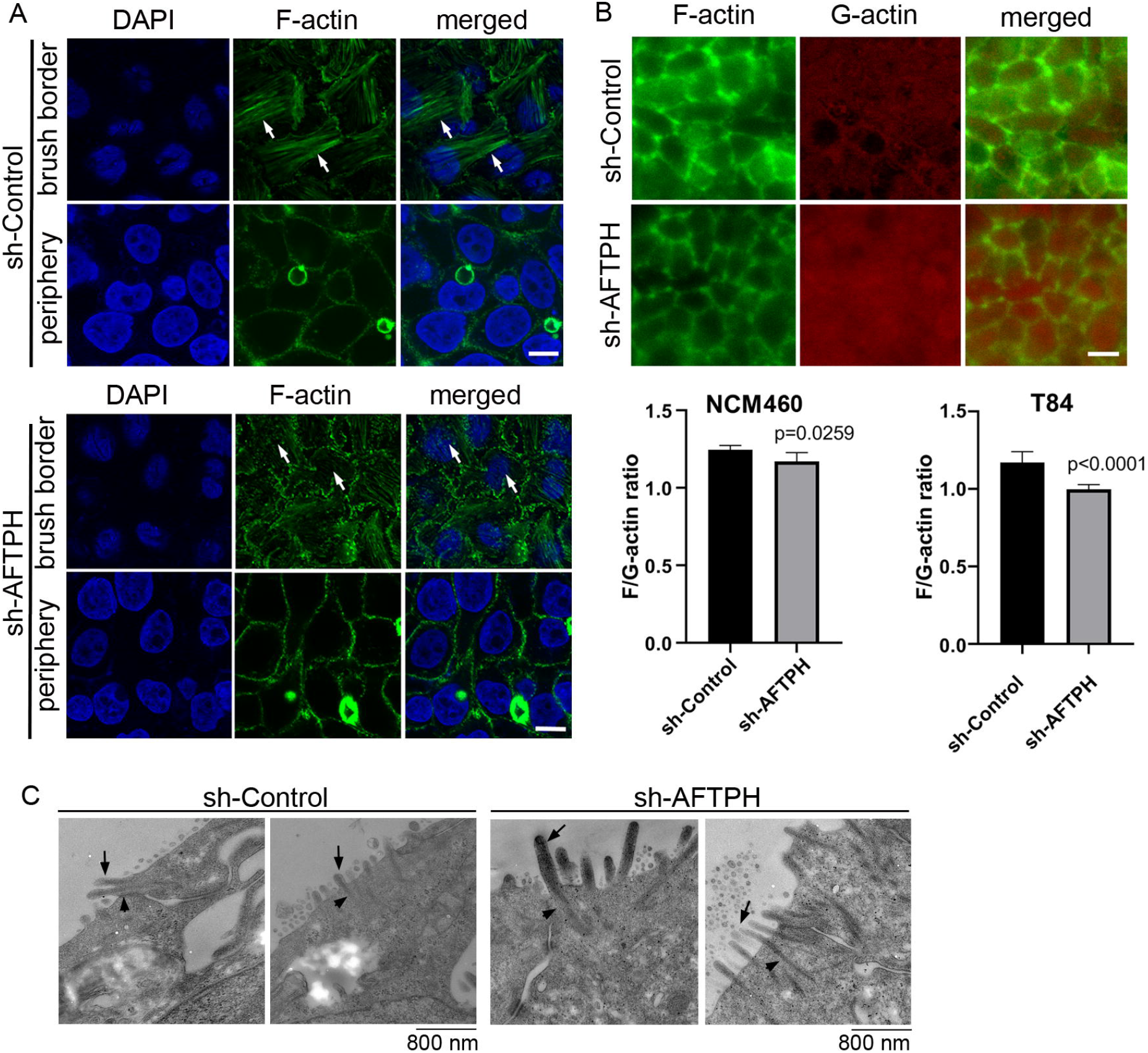
AFTPH depletion reduces actin polymerization and increases microvilli formation. (A) NCM460 sh-control and sh-AFTPH cells were seeded and fixed with 4% PFA. F-actin of at brush border and periphery were stained and visualized by confocal microscopy. This experiment was repeated 3 times. (Scale = 10 μm) (B) F-actin at peri-junctional actomyosin ring and G-actin monomer in polarized T84 sh-control and sh-AFTPH monolayers were stained and visualized by fluorescence microscopy. (Scale = 10 μm). NCM460 and T84 sh-control and sh-AFTPH cells were cultured in 96-well plates. F- and G-actin were stained and a fluriometer was used to measure their intensity. F/G-actin ratios were then calculated. Mean ± SEM. (n=10). (C) Polarized T84 sh-control and sh-AFTPH monolayers were cultured and fixed as described in the Methods. Electron microscopy showing microvilli (arrow) and F-actin core (arrowhead) present in polarized T84 sh-control and sh-AFTPH monolayers. (Scale = 800 nm)

## Discussion

Previous studies have demonstrated that AFTPH plays a role in exocytosis (26), intracellular trafficking (23) and receptor recycling (25) in various cell types. While our studies have demonstrated reduced AFTPH levels in colonic tissues during colonic inflammation, potentially promoting proinflammatory signaling in colonic epithelial cells (21); we examined the potential role and molecular mechanism of AFTPH in regulating colonic epithelial permeability *in vitro* in our present study. Our results demonstrate that reduced AFTPH levels in colonic epithelial cells led to impaired TEER, increased Na^+^ ion flow, and increased dextran permeability. Notably, our experimental evidence strongly suggested that differences in TEER between control and AFTPH-deficient colonic epithelial monolayers were largely due to differences in Na^+^ ion flow (Fig. 2). However, no candidates from the claudin family, including claudin-2, which can form Na^+^ ion channel [review in (2), (7)], and Na^+^ ion transporters were commonly dysregulated in both Caco2-BBe and T84 AFTPH-silenced monolayers (Fig. 3&4). On the other hand, results from confocal microscopy suggested a novel finding that MLCK, in which its kinase activity is essential to contraction of peri-junctional actomyosin ring (15), is colocalized with AFTPH in both T84 and Caco2-BBe cells intracellularly (Fig. 5). Furthermore, examination of the molecular alterations after AFTPH gene-silencing showed that MLCK activity was increased, while actin polymerization was reduced in AFTPH-deficient T84 cells (Fig. 5&6). As for limitations in this study, although our studies suggested that surface localization of TJ complex is not affected by AFTPH deficiency, the intracellular localization of TJ proteins could not be determined due to acetone fixation of the cells (36). Therefore, the trafficking of these TJ proteins remains to be studied.

In this study, our results demonstrate that AFTPH deficiency leads to increased MLCK activity (Fig. 5), which coincides with increased Na^+^ ion flux in the presence of a transepithelial Na^+^ ion gradient (Fig. 2B). Furthermore, inhibition of MLCK activity reverses increased intestinal epithelial permeability (TEER) caused by elevated MLCK activity *in vitro* (Fig. 5), similar to results observed in previous studies (15,34,35). We hypothesize that increased MLCK activation in AFTPH-depleted colonic epithelial monolayers promotes contraction of peri-junctional actomyosin ring (15) and/or destabilization of TJ protein complex (34); thus, in the presence of transepithelial Na^+^ gradient (Fig. 2C), increased number of Na^+^ ions passes through transcellular space across colonic epithelium and leads to reduction in Na^+^ ion dilution potential when compared to control epithelial monolayers. Interestingly, AFTPH deficiency-induced MLCK activation did not result in a uniform increase in transepithelial dextran flux in both Caco2-BBe and T84 monolayers (Fig. 2D-F). Previous studies have also shown that activation of MLCK may lead to differential regulation of ion and macromolecules flux under different physiological stimuli. The most well-studied examples are regulation of paracellular flux during Na^+^-glucose cotransport and stimulation from tumor necrosis factor (TNF) through MLCK activation. During Na^+^-glucose cotransport, sodium-dependent glucose transporter (SGLT), located at the brush-border, is activated and this transactivates MLCK in intestinal epithelial cells (18,37). Increased pMLC2 after loss of AFTPH present at peri-junctional actomyosin ring is the result from activation of MLCK (18), potentially induces contraction of peri-junctional actomyosin ring (15) and thus, increases paracellular space in intestinal epithelium. Together with activation Na^+^ ion transporters, SLC9A3 (NHE3) (19) and Na^+^/K^+^ transporting ATPase, a Na^+^ ion concentration gradient is generated across colonic epithelium, allowing increased flow of water, along with small solute (size: <4Å), across epithelium through paracellular space (38). In TNF-induced paracellular flux, TNF, together with other proinflammatory cytokines, increases intestinal epithelial permeability (14,39) through activation of MLCK (14). Interestingly, TNF-induced MLCK activation allows macromolecules flux (e.g. sizes of dextran: > 14Å) across intestinal epithelium (14,39) and is the major mechanism promoting TNF-induced diarrhea (40). The detailed mechanisms leading to these two different forms of MLCK-dependent paracellular flux are not well understood.

On the other hand, our findings on colocalization of MLCK and AFTPH provided novel insights on regulation of MLCK activity. There are 2 isoforms of MLCKs expressed in mammalian muscle and non-muscle cells, namely long MLCK isoform (220 kDa) and short MLCK isoform (130 kDa) (41). In biopsies taken from human small intestine, long MLCK isoform is localized at the cytoplasm adjacent to brush borders (42). In contrast, in polarized Caco2-BBe cells, long MLCK isoforms were localized in the cytoplasm, but was translocated to peri-junctional actomyosin ring upon TNF stimulation (42). Our results provided evidence supporting the notion that MLCK can be localized in cytosol with AFTPH, potentially at TGN or early endosomes (25) in polarized Caco2-BBe and T84 monolayers (Fig. 5). In addition, our novel findings on intracellular MLCK/AFTPH colocalization may have important implications in revealing mechanisms regulating of MLCK activities (Fig. 7A). Analysis of amino acid sequences suggests that both MLCK isoforms comprise of a catalytic domain, 2 actin-binding domains, 2 calmodulin-binding domains and 1 myosin-binding domain [review in (43), Fig. 7B]. This is evident in co-localization studies in fibroblasts revealing that the long MLCK isoform colocalizes with F-actin, while short MLCK isoform is localized with non-muscle myosin 2A (NMMIIA) (41,44). Non-muscle myosin 2 (NMMIIs) belong to myosin superfamily and are structurally grouped as heterohexamers [review in (45,46)]. As shown in Fig. 7B, NMMIIs are comprised of 2 heavy chains which possess enzymatic activities, which drive ATP-dependent actin filament movement; 2 essential light chains which stabilize the heavy chain structure; and regulatory light chain (also called MLC2) which is the target of MLCK catalytic domain and regulates NMMs activity in actomyosin complex as described (37,40,44,47). Previous studies demonstrated that the heavy chain of NMMIIs and AFTPH have similar functions in intracellular trafficking in polarized epithelial cells (22,24,25,48); while NMMII heterohexamer is a binding partner of AFTPH in HeLa cells (23). Therefore, our results highly suggest that AFTPH is essential in mediating the direct interaction between MLCK and NMMII/MLC2 and this interaction, subsequently, affect the integrity of intestinal barrier (Fig. 7).

**Fig. 7.**
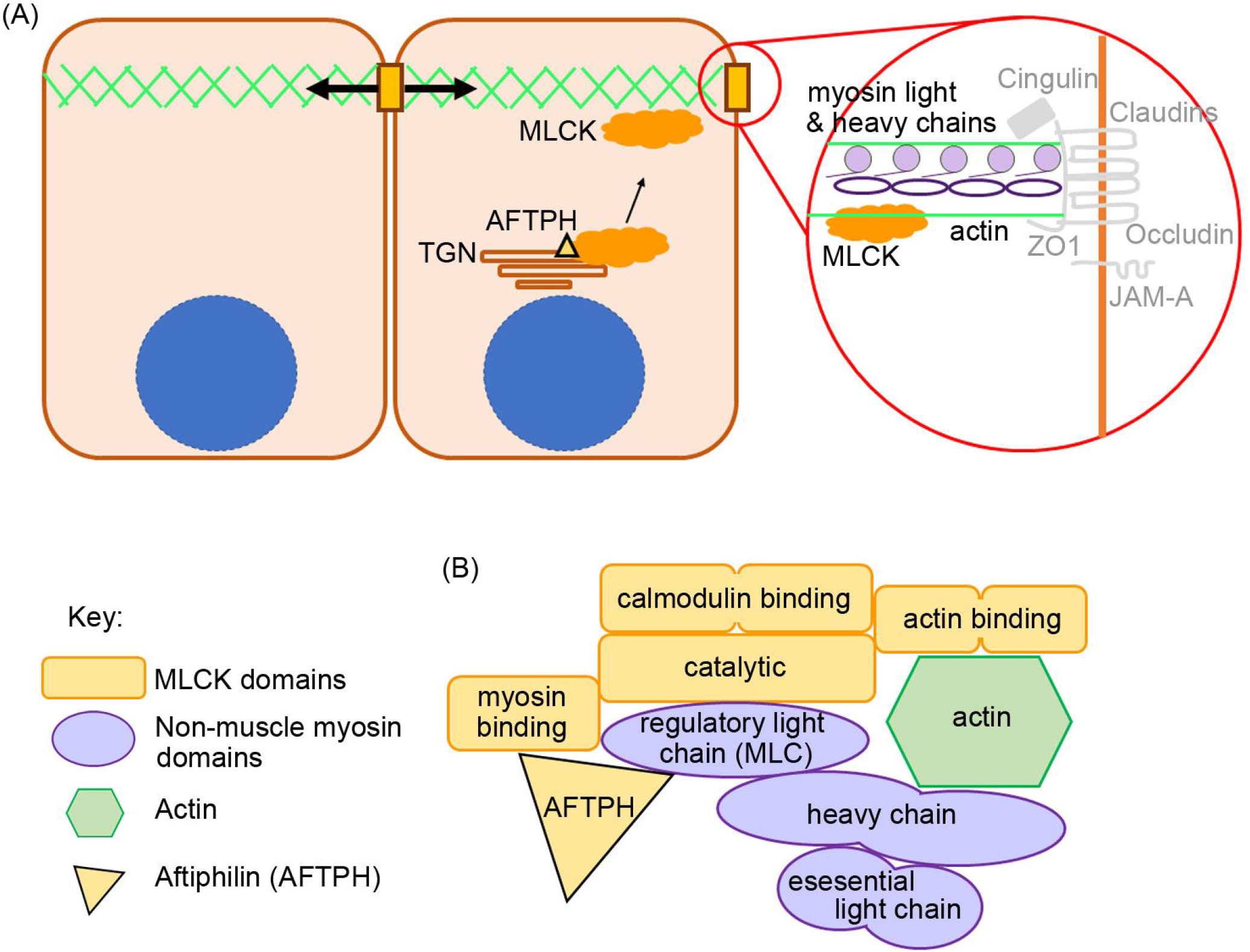
Proposed role of AFTPH in facilitating translocation of MLCK. (A) A diagram showing the proposed role of AFTPH in transporting MLCK from trans-golgi network (TGN) to peri-junctional actomyosin ring to promote the phosphorylation of MLC. (B) A diagram showing the proposed binding complex of AFTPH/ MLCK/non-muscle myosin.

There is an implication that the AFTPH / MLCK / NMMII network may also explain changes in Occludin expression in AFTPH-depleted cells. Similar to the results of the present study (Fig. 4), depletion of NMMIIA also results in functional defective, but structurally normal cellular junctions in intestinal epithelial monolayers (49). In particular, although localization of Occludin remains similar under NMMIIA depletion, upon destabilization of cellular junctions induced by calcium depletion, the majority of Occludin remains in cell-cell junctions in NMMIIA-depleted monolayers, while this phenomenon was not observed in control monolayers (49). Although the roles of Occludin in regulating intestinal epithelial permeability and tight junction structural integrity remains to be defined (9,29,50-52), the results of our study suggest that potential AFTPH-NMMIIA interaction may affect Occludin intracellular trafficking, which may have important implications to Occludin translocation and formation of TJ protein complex (53-55).

Lastly, in addition to dysregulated intestinal permeability, we also observed lower polymerization of actin monomers, disorganized F-actin filaments and increased number and length of microvilli in AFTPH-deficient intestinal monolayers (Fig. 6). A recent study by Chinowsky *et al*. demonstrated that NMMII is involved in actin polymerization/depolymerization and growth of epithelial microvilli (56). Thus, AFTPH-dependent actin polymerization and microvilli growth may also be associated with NMMII function in intestinal epithelial cells. In conclusion, our results highly suggest that AFTPH-MLCK interaction play an essential role in regulating the development and stability of cellular junctions and peri-junctional actomyosin ring in intestinal epithelial cells, potentially in collaboration with NMMIIA. Further investigation on the interactions between AFTPH, NMMIIA, MLCK may reveal novel insights to mechanisms regulating development and stability of cellular junctions in intestinal epithelium (Fig. 7).

## Experimental procedures

### Cell culture

Colonic epithelial NCM460 cells were maintained with M3D media (INCELL Inc.); Caco2-Bbe (ATCC CRL-2102) and T84 cells (ATCC CCL-248) were maintained with DMEM and DMEM/F:12. Base media was supplemented with 10% fetal bovine serum (FBS) and 1% Penicillin/Streptomycin (ThermoFisher Inc., Calrsbad, CA). Stable AFTPH knock-down clones were created by transducing NCM460, Caco2-BBe and T84 cells with recombinant lentivirus (Dharmacon ™ SMARTvector ™) expressing short hairpin RNAs (shRNAs) against AFTPH (Nbla10388, Horizon Discovery) and selected to create stable knock-down clones. Control clones were created by transducing the colonic epithelial cell lines with recombinant lentivirus expressing short hairpin with scramble sequence. Individual transduced clones from control shRNA and sh-AFTPH transduction were generated and levels of AFTPH were verified using PCR and western blot analysis. 2 control clones and 3 sh-AFTPH clones were used in subsequent experiments.

### Measurement of transepithelial electrical resistance

To measure transepithelial electrical resistance (TEER), Caco2-BBe cells (1×10^5^ cells) or T84 cells (1.5 × ×10^5^ cells) were seeded on semi-permeable supports (0.33 cm^2^, pore size: 0.4um Corning) and cultured for 4-7 days. A Millicell ERS-2 epithelial volt-ohm meter (Millipore) was used to measure TEER. In the experiments described below, Caco2-BBe and T84 cells carrying sh-control had a TEER of at least 400 mΩ·cm^2^ and 2000 mΩ·cm^2^, respectively.

For dilution potential experiments, cells were seeded and cultured on the semi-permeable supports for 4-7 days as described previously by Buchert et al. (57). In brief, background transepithelial voltage (TEV, P_bkgd_) was measured by recording the voltage difference between apical and basal chamber of a permeable support without cells. Culture medium was replaced with Buffer A (120 mM NaCl, 10 mM HEPES, pH 7.4, 5 mM KCl, 10 mM NaHCO_3_, 1.2 mM CaCl_2_, and 1 mM MgSO_4_) in both apical and basal chamber and TEV in Solution A (P_A_) was measured. Then, a second TEV measurement (P_B_) was performed immediately after Buffer A in apical chamber was replaced by Buffer B (60 mM NaCl, 120 mM mannitol, 10 mM HEPES, pH 7.4, 5 mM KCl, 10 mM NaHCO_3_, 1.2 mM CaCl_2_, and 1 mM MgSO_4_). Dilution potential (ΔP) was calculated as follows: (P_A_ -P_bkgd_) -P_B_.

To measure dextran permeability, dextran polymers tagged with Alexa Fluor™ 680 (3,000, 10,000 MW, Thermofisher) and dextran tagged with fluorescein (40,000 MW, Sigma Aldrich) were resuspended in 100 μl HBSS (4 mg/ml, 5 mg/ml, 25 mg/ml, respectively) and placed in the apical chamber of the Transwell inserts. Samples from basolateral chambers were collected at 1 and 2 hours after the start of the experiment and signals from the samples were read by a fluorimeter (BioTek Instruments, Winooski, VT, USA) to determine the amount of dextran passing through the epithelial monolayers.

### Filamentous-/ Globular (F/G) -actin ratio

Stable NCM460 and T84 cell lines carrying sh-AFTPH and sh-control RNAs were seeded in 96-well black-walled microplates for fluorescence-based assays (Molecular Probes®, ThermoFisher). The cells were fixed 24 hours after seeding in a 4% buffered formaldehyde solution, pH 6.9 (Millipore Sigma) for 20 minutes at room temperature. The fixed cells were then permeabilized with 0.1% Triton-X 100 in PBS and stained with ActinGreen 488 ReadyProbes and Dexyribonuclease 1, Alexa Fluor ™ 594 conjugate (ThermoFisher) at room temperature for 20 minutes. The stained cells were then washed with 0.1% Tween-20 in PBS. The signals from the cells were then detected by a fluorimeter (BioTek). F/G-actin ratio of individual wells were calculated as follows: OD488(cell) -OD488(background) / OD594(cell) – OD 594(background).

### Fluorescence microscopy

To visualize F-actin in NCM460 cells, cells were fixed and stained as described above, using ActinGreen 488 ReadyProbes. For TJ proteins, Caco2-BBe and T84 epithelial monolayers were cultured on permeable support and fixed with pre-cooled acetone for 20 minutes at -20°C. The cells were then washed with PBS and stained with antibodies against tight junction proteins, such as, JAM-A (SAB4200468, Millipore Sigma), Cingulin (117796, Abcam), AFTPH (122239, Abcam), Occludin (#71-1500, Invitrogen), MLCK (# PA5-79716, Invitrogen) overnight at 4°C. After washing with 0.1% Tween-20 in PBS, the cells were stained with appropriate secondary antibodies. To visualize actin monomers and filaments in NCM460 and polarized T84 cells, cells were fixed and stained as stated above. The localization of different tight junction proteins, actin filaments were imaged with a Zeiss LSM 510 Meta laser scanning confocal microscope using a Zeiss 63x Plan-Apo/1.4 oil immersion objective (numerical aperture 1.4).

### Transmission electronic microscopy

Polarized T84 monolayers were fixed in 2.5% glutaraldehyde and 2% formaldehyde in 0.1 M sodium phosphate buffer (PB) overnight at 4 °C. After washing, samples were post-fixed in 1% osmium tetroxide in 0.1M PB, and dehydrated through a graded series of ethanol concentrations. After infiltration with Eponate 12 resin, the samples were embedded in fresh Eponate 12 resin and polymerized at 60°C for 48 hours. Ultrathin sections of 77 nm thickness were prepared and placed on formvar carbon coated copper grids and stained with uranyl acetate and Reynolds’ lead citrate. The grids were examined using a JEOL 100CX transmission electron microscope at 60 kV and images were captured by an AMT digital camera (Advanced Microscopy Techniques Corporation, model XR611) at Electron Microscopy Core Facility, UCLA Brain Research Institute.

### Quantitative RT-PCR

Total RNA from Caco2-BBe epithelial monolayers was isolated using standard TRIzol reagent protocol (Life Technologies, Carlsbad, CA). A High Capacity cDNA Reverse Transcription Kit (ThermoFisher) used to generate cDNA libraries from equal amounts of total RNA (500 ng). Quantitative RT-PCR (qRT-PCR) for various gene expression was performed using iTaq Universal SYBR Green Mix (Bio-Rad) as described previously (58). Specific primers against CLDN1, CLDN2, CLDN3, CLDN4 and OCLDN were purchased from Integrated DNA Technologies. The sequence of the primers are listed in Table 1.

**Table 1.**
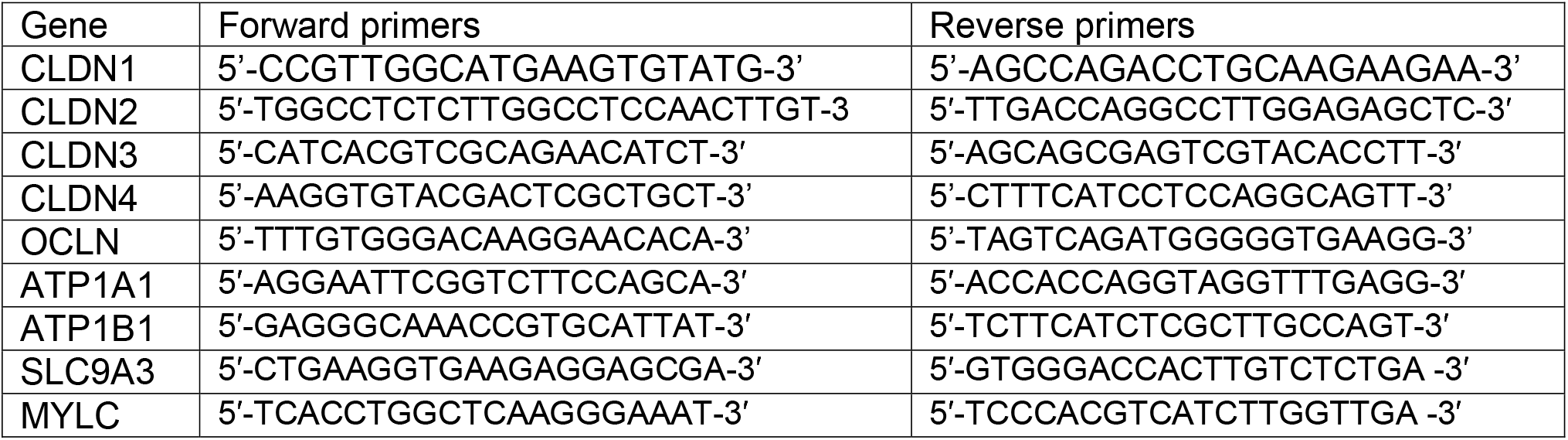

### Western Blots

Confluent T84 epithelial monolayers were lysed in RIPA buffer supplemented with protease inhibitors (ThermoFisher) and denatured at 95°C for 5 minutes. Samples were added to a 4% to 20% SDS-containing polyacrylamide gel (Bio-Rad Laboratories), and a Trans-Blot Turbo system (Bio-Rad Laboratories) was used to transfer proteins to PVDF membranes. Membranes were blocked (phosphate-buffered saline, 5% nonfat dry milk, 0.05% Tween-20) and probed with antibodies followed by corresponding horseradish peroxidase–labeled secondary antibodies. A ChemiDoc Touch Imager and Clarity enhanced chemiluminescence kit were used to develop the blots (Bio-Rad Laboratories). Antibodies used include Claudin 1 (SAB4200462, Sigma Aldrich), Claudin 2 (ab53032, Abcam), Claudin 3 (SAB4500435, Sigma Aldrich), Occludin (#71-1500, Invitrogen), pMLC2 (3671, Cell Signaling). Data are represented by cropped images from the original membranes.

### Statistical analysis

Unless otherwise stated, all experiments were performed three independent times. A Student *t-test* was used to compare the statistical significance between control and sh-AFTPH groups. Samples run in triplicate and data represent mean ± SD.

## Abbreviations

AFTPH: Aftiphilin
MLCK: myosin light chain kinase
TJ: tight junction
TEER: transepithelial electric resistance
TGN: trans-golgi network

## Data availability

The raw data from the manuscript is available upon request.

## Acknowledgments

We would like to thank the Imaging and Stem Cell Biology Core (ISCB, NIH/NIDDL Center Grant P30 DK 41301) and UCLA Brain Research Institute Electron Microscopy core facility that provided technical support for the confocal and transmission electron microscopy.

## References

1. Nusrat, A., Turner, J. R., and Madara, J. L. (2000) Molecular physiology and pathophysiology of tight junctions. IV. Regulation of tight junctions by extracellular stimuli: nutrients, cytokines, and immune cells. Am J Physiol Gastrointest Liver Physiol 279, G851–857

2. Garcia-Hernandez, V., Quiros, M., and Nusrat, A. (2017) Intestinal epithelial claudins: expression and regulation in homeostasis and inflammation. Ann N Y Acad Sci 1397, 66–79

3. Mrsny, R. J., Brown, G. T., Gerner-Smidt, K., Buret, A. G., Meddings, J. B., Quan, C., Koval, M., and Nusrat, A. (2008) A key claudin extracellular loop domain is critical for epithelial barrier integrity. Am J Pathol 172, 905–915

4. Bruewer, M., Luegering, A., Kucharzik, T., Parkos, C. A., Madara, J. L., Hopkins, A. M., and Nusrat, A. (2003) Proinflammatory cytokines disrupt epithelial barrier function by apoptosis-independent mechanisms. J Immunol 171, 6164–6172

5. Saeedi, B. J., Kao, D. J., Kitzenberg, D. A., Dobrinskikh, E., Schwisow, K. D., Masterson, J. C., Kendrick, A. A., Kelly, C. J., Bayless, A. J., Kominsky, D. J., Campbell, E. L., Kuhn, K. A., Furuta, G. T., Colgan, S. P., and Glover, L. E. (2015) HIF-dependent regulation of claudin-1 is central to intestinal epithelial tight junction integrity. Mol Biol Cell 26, 2252–2262

6. Rosenthal, R., Milatz, S., Krug, S. M., Oelrich, B., Schulzke, J. D., Amasheh, S., Günzel, D., and Fromm, M. (2010) Claudin-2, a component of the tight junction, forms a paracellular water channel. J Cell Sci 123, 1913–1921

7. Wada, M., Tamura, A., Takahashi, N., and Tsukita, S. (2013) Loss of claudins 2 and 15 from mice causes defects in paracellular Na+ flow and nutrient transport in gut and leads to death from malnutrition. Gastroenterology 144, 369–380

8. Raju, P., Shashikanth, N., Tsai, P. Y., Pongkorpsakol, P., Chanez-Paredes, S., Steinhagen, P. R., Kuo, W. T., Singh, G., Tsukita, S., and Turner, J. R. (2020) Inactivation of paracellular cation-selective claudin-2 channels attenuates immune-mediated experimental colitis in mice. J Clin Invest 130, 5197–5208

9. Al-Sadi, R., Khatib, K., Guo, S., Ye, D., Youssef, M., and Ma, T. (2011) Occludin regulates macromolecule flux across the intestinal epithelial tight junction barrier. Am J Physiol Gastrointest Liver Physiol 300, G1054–1064

10. Otani, T., Nguyen, T. P., Tokuda, S., Sugihara, K., Sugawara, T., Furuse, K., Miura, T., Ebnet, K., and Furuse, M. (2019) Claudins and JAM-A coordinately regulate tight junction formation and epithelial polarity. J Cell Biol 218, 3372–3396

11. Cereijido, M., Shoshani, L., and Contreras, R. G. (2000) Molecular physiology and pathophysiology of tight junctions. I. Biogenesis of tight junctions and epithelial polarity. Am J Physiol Gastrointest Liver Physiol 279, G477–482

12. Rodgers, L. S., and Fanning, A. S. (2011) Regulation of epithelial permeability by the actin cytoskeleton. Cytoskeleton (Hoboken) 68, 653–660

13. He, W. Q., Wang, J., Sheng, J. Y., Zha, J. M., Graham, W. V., and Turner, J. R. (2020) Contributions of Myosin Light Chain Kinase to Regulation of Epithelial Paracellular Permeability and Mucosal Homeostasis. Int J Mol Sci 21

14. Wang, F., Graham, W. V., Wang, Y., Witkowski, E. D., Schwarz, B. T., and Turner, J. R. (2005) Interferon-gamma and tumor necrosis factor-alpha synergize to induce intestinal epithelial barrier dysfunction by up-regulating myosin light chain kinase expression. Am J Pathol 166, 409–419

15. Shen, L., Black, E. D., Witkowski, E. D., Lencer, W. I., Guerriero, V., Schneeberger, E. E., and Turner, J. R. (2006) Myosin light chain phosphorylation regulates barrier function by remodeling tight junction structure. J Cell Sci 119, 2095–2106

16. Yu, D., Marchiando, A. M., Weber, C. R., Raleigh, D. R., Wang, Y., Shen, L., and Turner, J. R. (2010) MLCK-dependent exchange and actin binding region-dependent anchoring of ZO-1 regulate tight junction barrier function. Proc Natl Acad Sci U S A 107, 8237–8241

17. Magalhães, D., Cabral, J. M., Soares-da-Silva, P., and Magro, F. (2016) Role of epithelial ion transports in inflammatory bowel disease. Am J Physiol Gastrointest Liver Physiol 310, G460–476

18. Berglund, J. J., Riegler, M., Zolotarevsky, Y., Wenzl, E., and Turner, J. R. (2001) Regulation of human jejunal transmucosal resistance and MLC phosphorylation by Na(+)-glucose cotransport. Am J Physiol Gastrointest Liver Physiol 281, G1487–1493

19. Turner, J. R., Black, E. D., Ward, J., Tse, C. M., Uchwat, F. A., Alli, H. A., Donowitz, M., Madara, J. L., and Angle, J. M. (2000) Transepithelial resistance can be regulated by the intestinal brush-border Na(+)/H(+) exchanger NHE3. Am J Physiol Cell Physiol 279, C1918–1924

20. Bruewer, M., Samarin, S., and Nusrat, A. (2006) Inflammatory bowel disease and the apical junctional complex. Ann N Y Acad Sci 1072, 242–252

21. Law, I. K., Bakirtzi, K., Polytarchou, C., Oikonomopoulos, A., Hommes, D., Iliopoulos, D., and Pothoulakis, C. (2015) Neurotensin--regulated miR-133alpha is involved in proinflammatory signalling in human colonic epithelial cells and in experimental colitis. Gut 64, 1095–1104

22. Ikonen, E., de Almeid, J. B., Fath, K. R., Burgess, D. R., Ashman, K., Simons, K., and Stow, J. L. (1997) Myosin II is associated with Golgi membranes: identification of p200 as nonmuscle myosin II on Golgi-derived vesicles. J Cell Sci 110 (Pt 18), 2155–2164

23. Hirst, J., Borner, G. H., Harbour, M., and Robinson, M. S. (2005) The aftiphilin/p200/gamma-synergin complex. Mol Biol Cell 16, 2554–2565

24. Burman, J. L., Wasiak, S., Ritter, B., de Heuvel, E., and McPherson, P. S. (2005) Aftiphilin is a component of the clathrin machinery in neurons. FEBS Lett 579, 2177–2184

25. Law, I. K., Jensen, D., Bunnett, N. W., and Pothoulakis, C. (2016) Neurotensin-induced miR-133alpha expression regulates neurotensin receptor 1 recycling through its downstream target aftiphilin. Sci Rep 6, 22195

26. Lui-Roberts, W. W., Ferraro, F., Nightingale, T. D., and Cutler, D. F. (2008) Aftiphilin and gamma-synergin are required for secretagogue sensitivity of Weibel-Palade bodies in endothelial cells. Mol Biol Cell 19, 5072–5081

27. Moyer, M. P., Manzano, L. A., Merriman, R. L., Stauffer, J. S., and Tanzer, L. R. (1996) NCM460, a normal human colon mucosal epithelial cell line. In Vitro Cell Dev Biol Anim 32, 315–317

28. Devriese, S., Van den Bossche, L., Van Welden, S., Holvoet, T., Pinheiro, I., Hindryckx, P., De Vos, M., and Laukens, D. (2017) T84 monolayers are superior to Caco-2 as a model system of colonocytes. Histochem Cell Biol 148, 85–93

29. Buschmann, M. M., Shen, L., Rajapakse, H., Raleigh, D. R., Wang, Y., Lingaraju, A., Zha, J., Abbott, E., McAuley, E. M., Breskin, L. A., Wu, L., Anderson, K., Turner, J. R., and Weber, C. R. (2013) Occludin OCEL-domain interactions are required for maintenance and regulation of the tight junction barrier to macromolecular flux. Mol Biol Cell 24, 3056–3068

30. Laukoetter, M. G., Nava, P., Lee, W. Y., Severson, E. A., Capaldo, C. T., Babbin, B. A., Williams, I. R., Koval, M., Peatman, E., Campbell, J. A., Dermody, T. S., Nusrat, A., and Parkos, C. A. (2007) JAM-A regulates permeability and inflammation in the intestine in vivo. J Exp Med 204, 3067–3076

31. Bazzoni, G., Martinez-Estrada, O. M., Orsenigo, F., Cordenonsi, M., Citi, S., and Dejana, E. (2000) Interaction of junctional adhesion molecule with the tight junction components ZO-1, cingulin, and occludin. J Biol Chem 275, 20520–20526

32. Cordenonsi, M., D’Atri, F., Hammar, E., Parry, D. A., Kendrick-Jones, J., Shore, D., and Citi, S. (1999) Cingulin contains globular and coiled-coil domains and interacts with ZO-1, ZO-2, ZO-3, and myosin. J Cell Biol 147, 1569–1582

33. Citi, S., Sabanay, H., Jakes, R., Geiger, B., and Kendrick-Jones, J. (1988) Cingulin, a new peripheral component of tight junctions. Nature 333, 272–276

34. Jin, Y., and Blikslager, A. T. (2016) Myosin light chain kinase mediates intestinal barrier dysfunction via occludin endocytosis during anoxia/reoxygenation injury. Am J Physiol Cell Physiol 311, C996–C1004

35. Feighery, L. M., Cochrane, S. W., Quinn, T., Baird, A. W., O’Toole, D., Owens, S. E., O’Donoghue, D., Mrsny, R. J., and Brayden, D. J. (2008) Myosin light chain kinase inhibition: correction of increased intestinal epithelial permeability in vitro. Pharm Res 25, 1377–1386

36. Hoetelmans, R. W., Prins, F. A., Cornelese-ten Velde, I., van der Meer, J., van de Velde, C. J., and van Dierendonck, J. H. (2001) Effects of acetone, methanol, or paraformaldehyde on cellular structure, visualized by reflection contrast microscopy and transmission and scanning electron microscopy. Appl Immunohistochem Mol Morphol 9, 346–351

37. Turner, J. R., Rill, B. K., Carlson, S. L., Carnes, D., Kerner, R., Mrsny, R. J., and Madara, J. L. (1997) Physiological regulation of epithelial tight junctions is associated with myosin light-chain phosphorylation. Am J Physiol 273, C1378–1385

38. Turner, J. R., Cohen, D. E., Mrsny, R. J., and Madara, J. L. (2000) Noninvasive in vivo analysis of human small intestinal paracellular absorption: regulation by Na+-glucose cotransport. Dig Dis Sci 45, 2122–2126

39. Ma, T. Y., Iwamoto, G. K., Hoa, N. T., Akotia, V., Pedram, A., Boivin, M. A., and Said, H. M. (2004) TNF-alpha-induced increase in intestinal epithelial tight junction permeability requires NF-kappa B activation. Am J Physiol Gastrointest Liver Physiol 286, G367–376

40. Clayburgh, D. R., Barrett, T. A., Tang, Y., Meddings, J. B., Van Eldik, L. J., Watterson, D. M., Clarke, L. L., Mrsny, R. J., and Turner, J. R. (2005) Epithelial myosin light chain kinase-dependent barrier dysfunction mediates T cell activation-induced diarrhea in vivo. J Clin Invest 115, 2702–2715

41. Blue, E. K., Goeckeler, Z. M., Jin, Y., Hou, L., Dixon, S. A., Herring, B. P., Wysolmerski, R. B., and Gallagher, P. J. (2002) 220- and 130-kDa MLCKs have distinct tissue distributions and intracellular localization patterns. Am J Physiol Cell Physiol 282, C451–460

42. Graham, W. V., He, W., Marchiando, A. M., Zha, J., Singh, G., Li, H. S., Biswas, A., Ong, M. L. D. M., Jiang, Z. H., Choi, W., Zuccola, H., Wang, Y., Griffith, J., Wu, J., Rosenberg, H. J., Snapper, S. B., Ostrov, D., Meredith, S. C., Miller, L. W., and Turner, J. R. (2019) Intracellular MLCK1 diversion reverses barrier loss to restore mucosal homeostasis. Nat Med 25, 690–700

43. Gao, Y., Ye, L. H., Kishi, H., Okagaki, T., Samizo, K., Nakamura, A., and Kohama, K. (2001) Myosin light chain kinase as a multifunctional regulatory protein of smooth muscle contraction. IUBMB Life 51, 337–344

44. Kassianidou, E., Hughes, J. H., and Kumar, S. (2017) Activation of ROCK and MLCK tunes regional stress fiber formation and mechanics via preferential myosin light chain phosphorylation. Mol Biol Cell 28, 3832–3843

45. De La Cruz, E.M., and Ostap, E. M. (2004) Relating biochemistry and function in the myosin superfamily. Curr Opin Cell Biol 16, 61–67

46. Krendel, M., and Mooseker, M. S. (2005) Myosins: tails (and heads) of functional diversity. Physiology (Bethesda) 20, 239–251

47. Wendt, T., Taylor, D., Trybus, K. M., and Taylor, K. (2001) Three-dimensional image reconstruction of dephosphorylated smooth muscle heavy meromyosin reveals asymmetry in the interaction between myosin heads and placement of subfragment 2. Proc Natl Acad Sci U S A 98, 4361–4366

48. Müsch, A., Cohen, D., and Rodriguez-Boulan, E. (1997) Myosin II is involved in the production of constitutive transport vesicles from the TGN. J Cell Biol 138, 291–306

49. Ivanov, A. I., Bachar, M., Babbin, B. A., Adelstein, R. S., Nusrat, A., and Parkos, C. A. (2007) A unique role for nonmuscle myosin heavy chain IIA in regulation of epithelial apical junctions. PLoS One 2, e658

50. Yu, A. S., McCarthy, K. M., Francis, S. A., McCormack, J. M., Lai, J., Rogers, R. A., Lynch, R. D., and Schneeberger, E. E. (2005) Knockdown of occludin expression leads to diverse phenotypic alterations in epithelial cells. Am J Physiol Cell Physiol 288, C1231–1241

51. Saitou, M., Furuse, M., Sasaki, H., Schulzke, J. D., Fromm, M., Takano, H., Noda, T., and Tsukita, S. (2000) Complex phenotype of mice lacking occludin, a component of tight junction strands. Mol Biol Cell 11, 4131–4142

52. Schulzke, J. D., Gitter, A. H., Mankertz, J., Spiegel, S., Seidler, U., Amasheh, S., Saitou, M., Tsukita, S., and Fromm, M. (2005) Epithelial transport and barrier function in occludin-deficient mice. Biochim Biophys Acta 1669, 34–42

53. Murakami, T., Felinski, E. A., and Antonetti, D. A. (2009) Occludin phosphorylation and ubiquitination regulate tight junction trafficking and vascular endothelial growth factor-induced permeability. J Biol Chem 284, 21036–21046

54. Raleigh, D. R., Boe, D. M., Yu, D., Weber, C. R., Marchiando, A. M., Bradford, E. M., Wang, Y., Wu, L., Schneeberger, E. E., Shen, L., and Turner, J. R. (2011) Occludin S408 phosphorylation regulates tight junction protein interactions and barrier function. J Cell Biol 193, 565–582

55. Bolinger, M. T., Ramshekar, A., Waldschmidt, H. V., Larsen, S. D., Bewley, M. C., Flanagan, J. M., and Antonetti, D. A. (2016) Occludin S471 Phosphorylation Contributes to Epithelial Monolayer Maturation. Mol Cell Biol 36, 2051–2066

56. Chinowsky, C. R., Pinette, J. A., Meenderink, L. M., Lau, K. S., and Tyska, M. J. (2020) Nonmuscle myosin-2 contractility-dependent actin turnover limits the length of epithelial microvilli. Mol Biol Cell 31, 2803–2815

57. Buchert, M., Turksen, K., and Hollande, F. (2012) Methods to examine tight junction physiology in cancer stem cells: TEER, paracellular permeability, and dilution potential measurements. Stem Cell Rev Rep 8, 1030–1034

58. Soroosh, A., Rankin, C. R., Polytarchou, C., Lokhandwala, Z. A., Patel, A., Chang, L., Pothoulakis, C., Iliopoulos, D., and Padua, D. M. (2019) miR-24 Is Elevated in Ulcerative Colitis Patients and Regulates Intestinal Epithelial Barrier Function. Am J Pathol 189, 1763–1774

